# Cevipabulin-tubulin complex reveals a novel agent binding site on α-tubulin with tubulin degradation effect

**DOI:** 10.1101/2020.09.11.293563

**Authors:** Jianhong Yang, Yamei Yu, Yong Li, Wei Yan, Haoyu Ye, Lu Niu, Minghai Tang, Zhoufeng Wang, Zhuang Yang, Heying Pei, Haoche Wei, Min Zhao, Jiaolin Wen, Linyu Yang, Liang Ouyang, Yuquan Wei, Qiang Chen, Weimin Li, Lijuan Chen

## Abstract

Microtubule, composed of αβ-tubulin heterodimers, remains as one of the most popular anticancer targets for decades. To date, anti-microtubule drugs are mainly functionally divided into microtubule-destabilizing and microtubule-stabilizing agents while microtubule- or tubulin-degradation agents are rarely reported. Six known binding sites on tubulin dimer are identified with five sites on β-tubulin and only one site on α-tubulin (pironetin site), hinting compounds binding to α-tubulin are less well characterized. Cevipabulin, a microtubule-active antitumor clinical candidate, is widely accepted as a microtubule-stabilizing agent by binding to the vinblastine site. Our X-ray crystallography study reveals that, in addition binding to the vinblastine site, cevipabulin also binds to a novel site on α-tubulin (named the seventh site) which located near the nonexchangeable GTP. Interestingly, we find the binding of cevipabulin to the seventh site induces tubulin degradation. As the non-exchangeable GTP has structural role and is important for the stability of tubulin dimers, we propose and confirm the tubulin degradation mechanism as: Cevipabulin at the seventh site puts the αT5 loop outward to make the non-exchangeable GTP exchangeable, which reduces the stability of tubulin and results in its destabilization and degradation. Our results confirm a novel agent binding site on α-tubulin and shed light on the development of tubulin degraders as a new generation of anti-microtubule drugs targeting this novel site.

## Introduction

Microtubules play key roles in many important cell events, especially cell division, and thus remain as one of the most popular anticancer targets for decades [1, 2]. Microtubules are composed of αβ-tubulin heterodimers assembled into linear protofilaments and their packaging demands both lateral and longitudinal interactions between tubulins [3]. To date, various tubulin inhibitors have been reported to alter the lateral and/or longitudinal interactions to promote microtubule assembly or disassembly, including the clinical most popular anticancer drugs: vinca alkaloids, taxanes, eribulin *et al* [4, 5]. These drugs all target β-tubulin which has five different binding sites (colchicine, vinblastine, paclitaxel, laulimalide and maytansine sites) [5]. By overexpression of β-tubulin isoforms, especially βIII-tubulin, cancer cells are prone to become resistant to these therapies [6]. So far, the pironetin site is the only one located on α-tubulin [5, 7]. However, this site is too small and pironetin has six chiral centers in its molecular structure, making it difficult to be synthetized. Since the crystal structure of tubulin-pironetin was reported in 2016 [5, 7], no significant progress has been made in the design of pironetin-binding-site inhibitors or even analogues of pironetin.

Tubulin inhibitors are functionally divided into two categories: microtubule stabilization agents (MSAs) and microtubule destabilization agents (MDAs). MSAs that promote microtubule polymerization include the paclitaxel and laulimalide site inhibitors and structure biological studies reveal they both stabilize the M-loop to enhance lateral interactions to promote tubulin polymerization [8, 9]. MDAs that inhibit microtubule polymerization contain the colchicine, vinblastine, maytansine and pironetin site inhibitors. Colchicine binds to intra-dimer interfaces to prevent tubulin dimers from adopting a “straight conformation” thus inhibiting lateral interactions [10]. Maytansine and pironetin bind to the inter-dimer interfaces to inhibit longitudinal interactions [5, 7, 11] and vinblastine also binds to the inter-dimer interfaces however acts as a wedge to enhance abnormal longitudinal interactions finally self-associate into spiral aggregates [3]. Recently, some tubulin degradation agents were reported such as T0070907, T007-1 and withaferin A [12, 13], while the degradation mechanism was unclear.

Cevipabulin (or TTI-237) is a synthetic tubulin inhibitor with *in vivo* anticancer activity and has been used in clinical trials investigating the treatment of advanced malignant solid tumors [14]. Competition experiment showed it competed with ^3^H-vinblastine but not ^3^H-paclitaxel for binding to microtubules, indicating it binds to the classic tubulin-depolymerization vinblastine site [15]. However, an *in vitro* tubulin polymerization assay exhibited that cevipabulin did not inhibit tubulin polymerization as vinblastine but promoted tubulin polymerization as paclitaxel [15]. These studies concluded that cevipabulin seems displaying mixed properties between paclitaxel and vinblastine. Kovalevich *et al*. has found that cevipabulin could promote tubulin degradation [16]. However, the degradation mechanism was not fully elucidated. Recently, Saez-Calvo *et al*. synthetized an analogue of cevipabulin (named compound **2** in this paper) and got the crystal structure of compound **2-**tubulin complex (PDB code: 5NJH) and proved compound **2** binds to the vinblastine site of β-tubulin to enhance longitudinal interactions and induces the formation of tubulin bundles in cell, which further confirmed that compound **2** binding to vinblastine site to induce tubulin polymerization in a paclitaxel-like manner[17].

In this study, we further investigate the tubulin-inhibition mechanism of cevipabulin. The crystal structure of cevipabulin-tubulin complex reveals cevipabulin simultaneously binds to two spatially independent sites: the vinblastine site and a new site on α-tubulin (called the seventh site). Biochemical experiments results confirm the binding of cevipabulin to the novel site is responsible for its tubulin degradation effect. Our study reveals a novel binding site on α-tubulin related to tubulin degradation effect and lays a foundation for the rational design of new generation of anticancer drugs.

## Results

### Cevipabulin induces tubulin-heterodimer degradation

Cevipabulin was found to downregulate tubulin protein level in different cells[16]. To elucidate the cellular effect of cevipabulin at an early time point, we carried out label-free quantitative proteomic analysis on six-hour cevipabulin treated human cervical adenocarcinoma cell line HeLa. Cevipabulin significantly down-regulated the protein level of α, β-tubulin and their isoforms with high selectivity (Fig.1A). Immunoblotting study confirmed cevipabulin decreased tubulin proteins in HeLa, human colon colorectal carcinoma cell line Hct116, human large cell lung carcinoma cell line H460 and human B cell lymphoma cell SU-DHL-6 in a dose-dependent manner (Fig.1B) and time-dependent manner in HeLa cells (Fig. 1C), demonstrating that the reduction of tubulin was a common biochemical consequence of cevipabulin treatment in cancer cells. The quantitative PCR assay showed that cevipabulin had no effect on *α-* and *β-tubulin* mRNA levels (Fig.1D), indicating that the downregulation of tubulin protein by cevipabulin was post-transcriptional. MG132, a proteasome inhibitor, could completely block cevipabulin-induced tubulin degradation (Fig.1E). All these proved that cevipabulin promoted tubulin degradation in a proteasome-dependent pathway.

**Figure 1.**
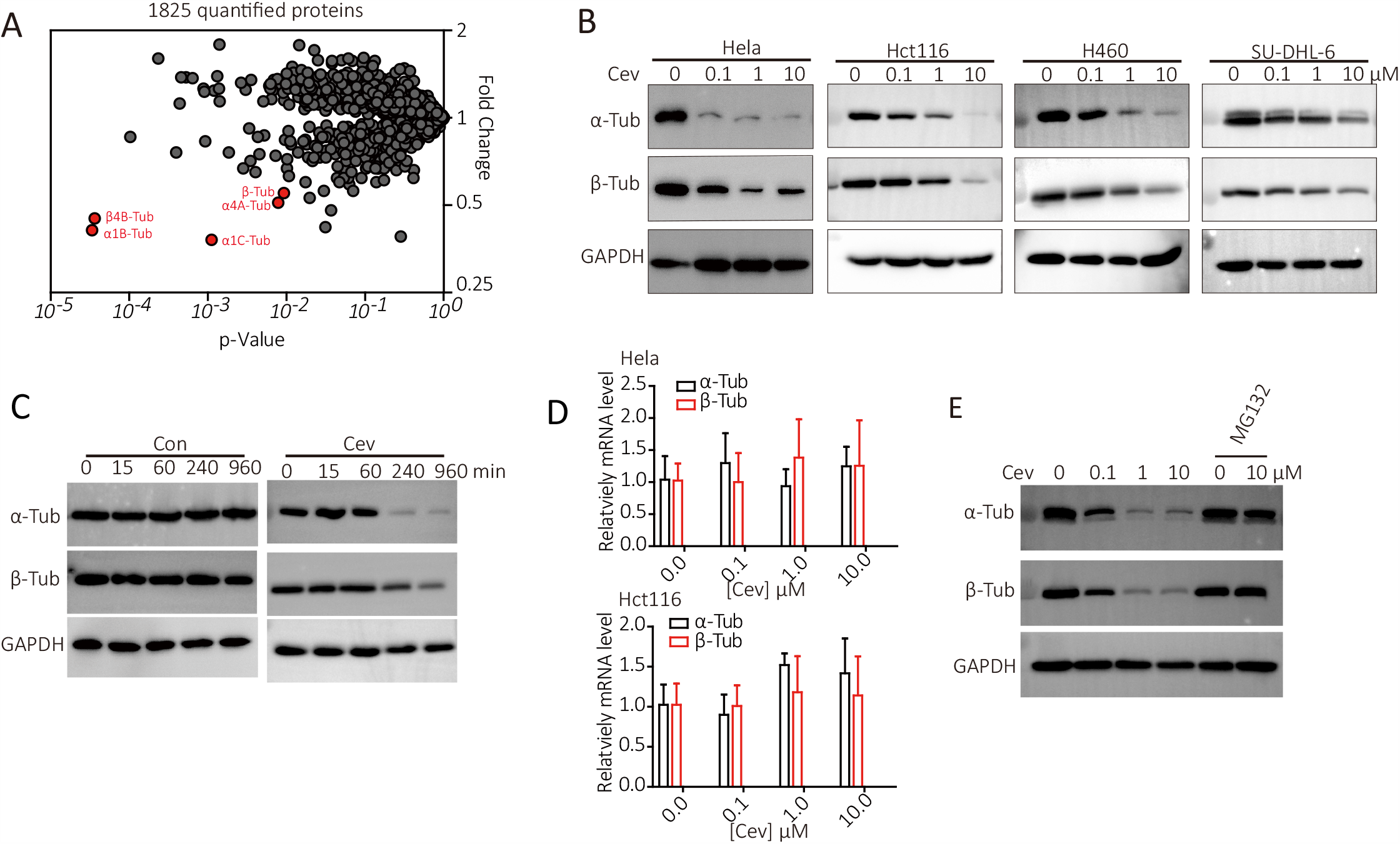
Cevipabulin promotes tubulin dimer degradation. (**A**) Label-free quantitative proteomic analysis of total proteins from HeLa cells treated with 1 μM cevipabulin for 6 h. This graph presents fold-changes of 1825 quantified proteins between cevipabulin and DMSO treatment groups versus the p value (t test; triplicate analysis). Three biological repetitions are performed. (**B**) Immunoblotting analysis of both α and β-tubulin levels in HeLa, Hct116, H460 and SU-DHL-6 cells, which all are treated with indicated concentrations of cevipabulin for 16 h. Results are representative of three independent experiments. (**C**) HeLa cells were treated with 1 μM cevipabulin for the indicated times and then the α and β-tubulin levels were detected by immunoblotting. Results are representative of two independent experiments. (**D**) HeLa and Hct116 cells were treated with indicated concentrations of cevipabulin for 16 hours, and then mRNA levels of both *α-tubulin* and *β-tubulin* were measured by quantitative-PCR. Data were shown as means ± SD of three independent experiments. **(E**) Cells were treated with or without MG132 (20 μM) for one hour before treated with different concentrations of cevipabulin for 16 hours. Protein levels of both α- and β-tubulin were detected by immunoblotting. Results are representative of two independent experiments. Cev: cevipabulin; Tub:tubulin.

### Crystal structure of cevipabulin-tubulin reveals its simultaneously binding to the vinblastine site and a novel site on α-tubulin

Previous studies concluded cevipabulin was an MSA binding to the vinblastine site [14, 18]. However, the detail interaction between tubulin and cevipabulin was not elucidated. To analyze the binding details of cevipabulin (Fig. 2A) to tubulin, we soaked cevipabulin into the crystals consisting of two tubulin heterodimers, one stathmin-like protein RB3 and one tubulin tyrosine ligase (T2R-TTL) [8]. The crystal structure of cevipabulin-tubulin complex was determined to be 2.6 Å resolution (Table 1). The whole structure was identical to that previously reported [8], in which two tubulin heterodimers were arranged in a head to tail manner (α1β1-α2β2) with the long helix RB3 comprising both dimers and tubulin tyrosine ligase docking onto α1-tubulin (Fig. 2B). The Fo–Fc difference electron density unambiguously revealed two cevipabulin molecules binding to two different sites (Fig. 2C and 2D): one at the inter-dimer interfaces between the β1- and α2-tubulin subunits (the vinblastine site) and the other one at the intra-dimer interfaces between α2- and β2-tubulin subunits (Fig. 2B) and the later binding region is a new binding site (here named as the seventh site).

**Table 1.**
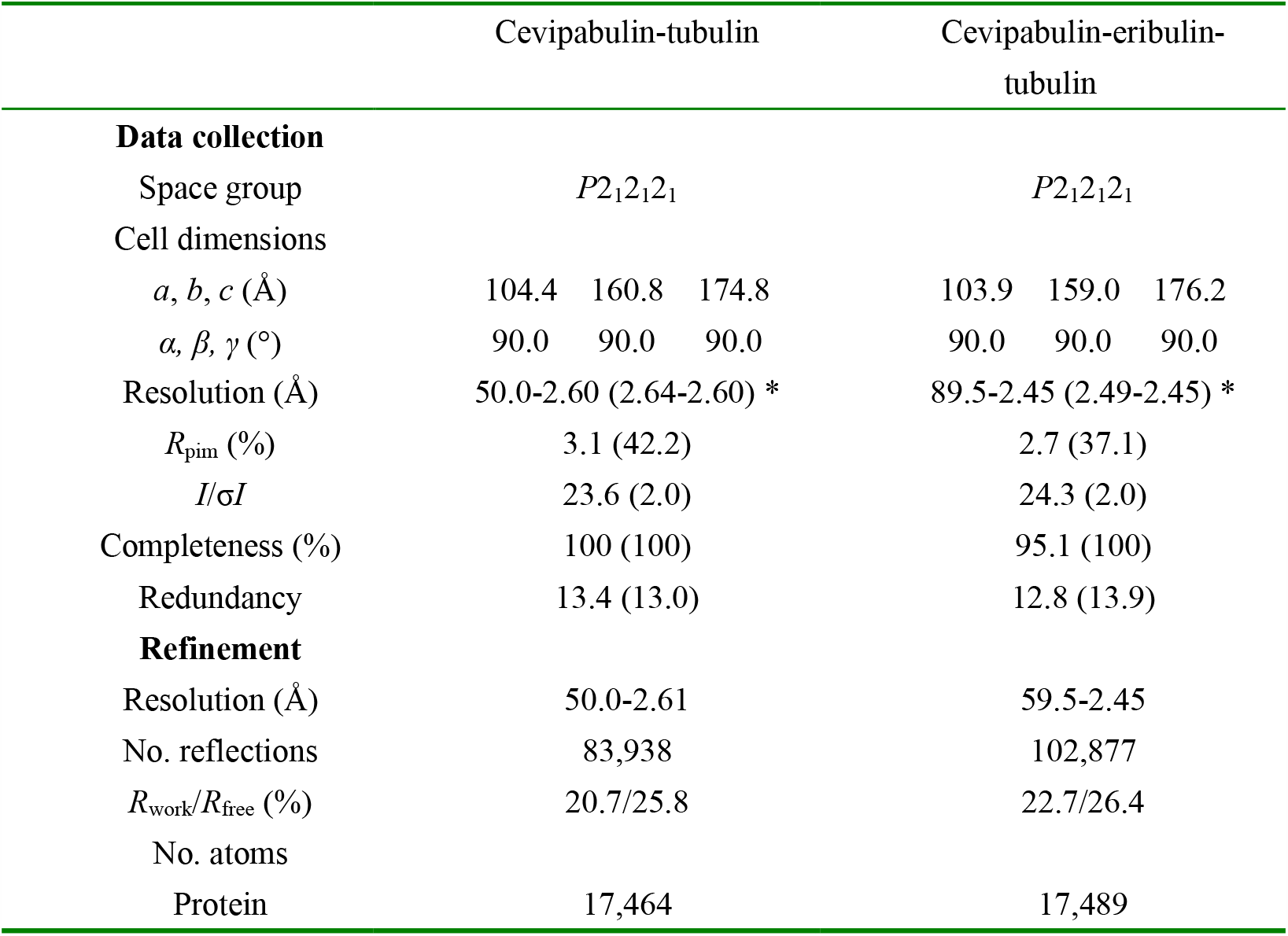
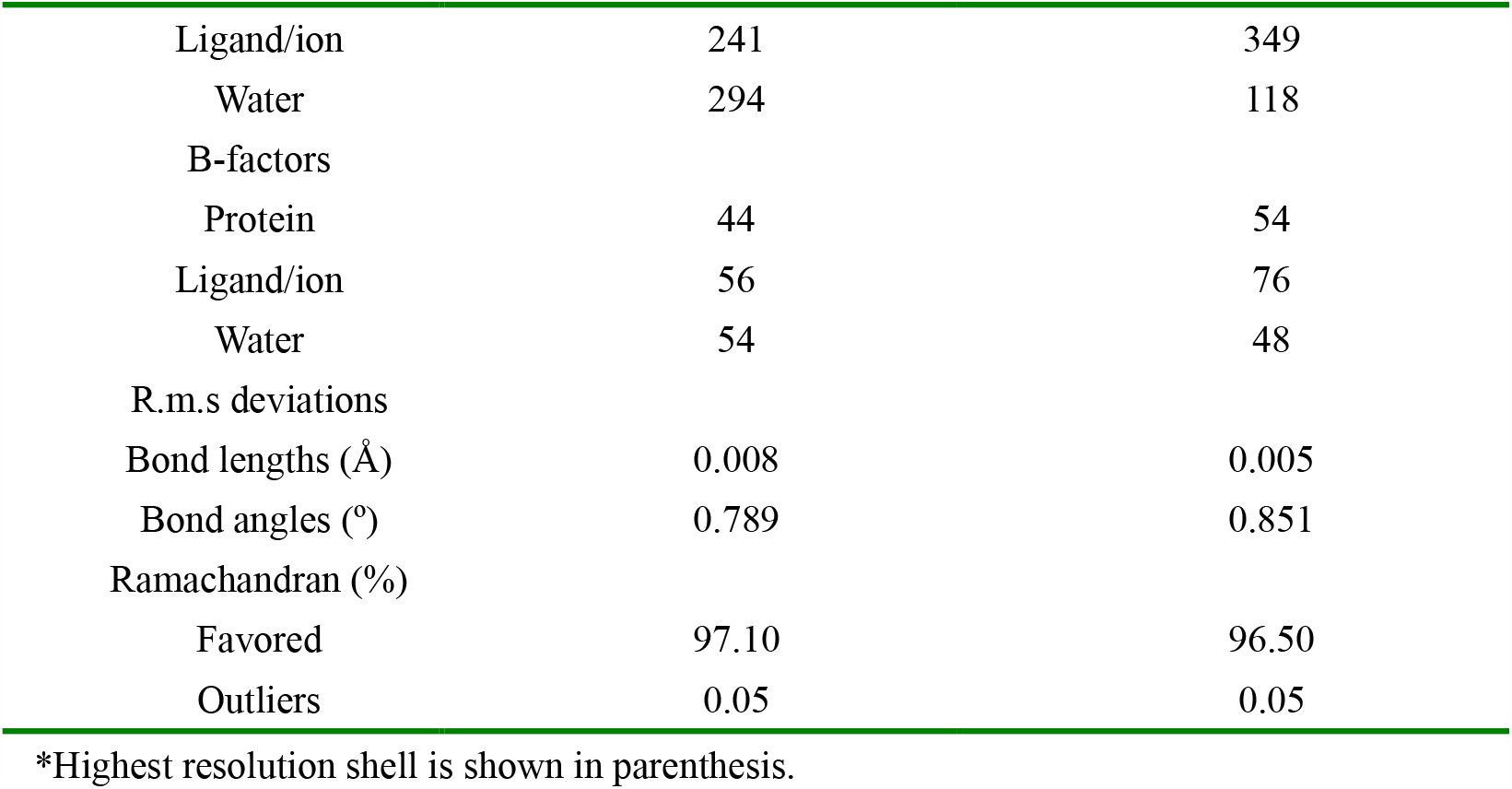
Data collection and refinement statistics.

|  | Cevipabulin-tubulin |  |  | Cevipabulin-eribulin-tubulin |  |  |
| --- | --- | --- | --- | --- | --- | --- |
| **Data collection** |  |  |  |  |  |  |
| Space group | $P2_12_12_1$ | | | $P2_12_12_1$ | | |
| Cell dimensions |  |  |  |  |  |  |
| $a, b, c$ (Å) | 104.4 | 160.8 | 174.8 | 103.9 | 159.0 | 176.2 |
| $\alpha, \beta, \gamma$ (°) | 90.0 | 90.0 | 90.0 | 90.0 | 90.0 | 90.0 |
| Resolution (Å) | 50.0-2.60 (2.64-2.60) * |  |  | 89.5-2.45 (2.49-2.45) * |  |  |
| $R_{\text{pim}}$ (%) | 3.1 (42.2) | | | 2.7 (37.1) | | |
| $I/\sigma I$ | 23.6 (2.0) | | | 24.3 (2.0) | | |
| Completeness (%) | 100 (100) |  |  | 95.1 (100) |  |  |
| Redundancy | 13.4 (13.0) |  |  | 12.8 (13.9) |  |  |
| **Refinement** |  |  |  |  |  |  |
| Resolution (Å) | 50.0-2.61 |  |  | 59.5-2.45 |  |  |
| No. reflections | 83,938 |  |  | 102,877 |  |  |
| $R_{\text{work}}/R_{\text{free}}$ (%) | 20.7/25.8 | | | 22.7/26.4 | | |
| No. atoms |  |  |  |  |  |  |
| Protein | 17,464 |  |  | 17,489 |  |  |

|  |  |  |
| --- | --- | --- |
| Ligand/ion | 241 | 349 |
| Water | 294 | 118 |
| B-factors |  |  |
| Protein | 44 | 54 |
| Ligand/ion | 56 | 76 |
| Water | 54 | 48 |
| R.m.s deviations |  |  |
| Bond lengths (Å) | 0.008 | 0.005 |
| Bond angles (°) | 0.789 | 0.851 |
| Ramachandran (%) |  |  |
| Favored | 97.10 | 96.50 |
| Outliers | 0.05 | 0.05 |

\*Highest resolution shell is shown in parenthesis.

**Figure 2.**
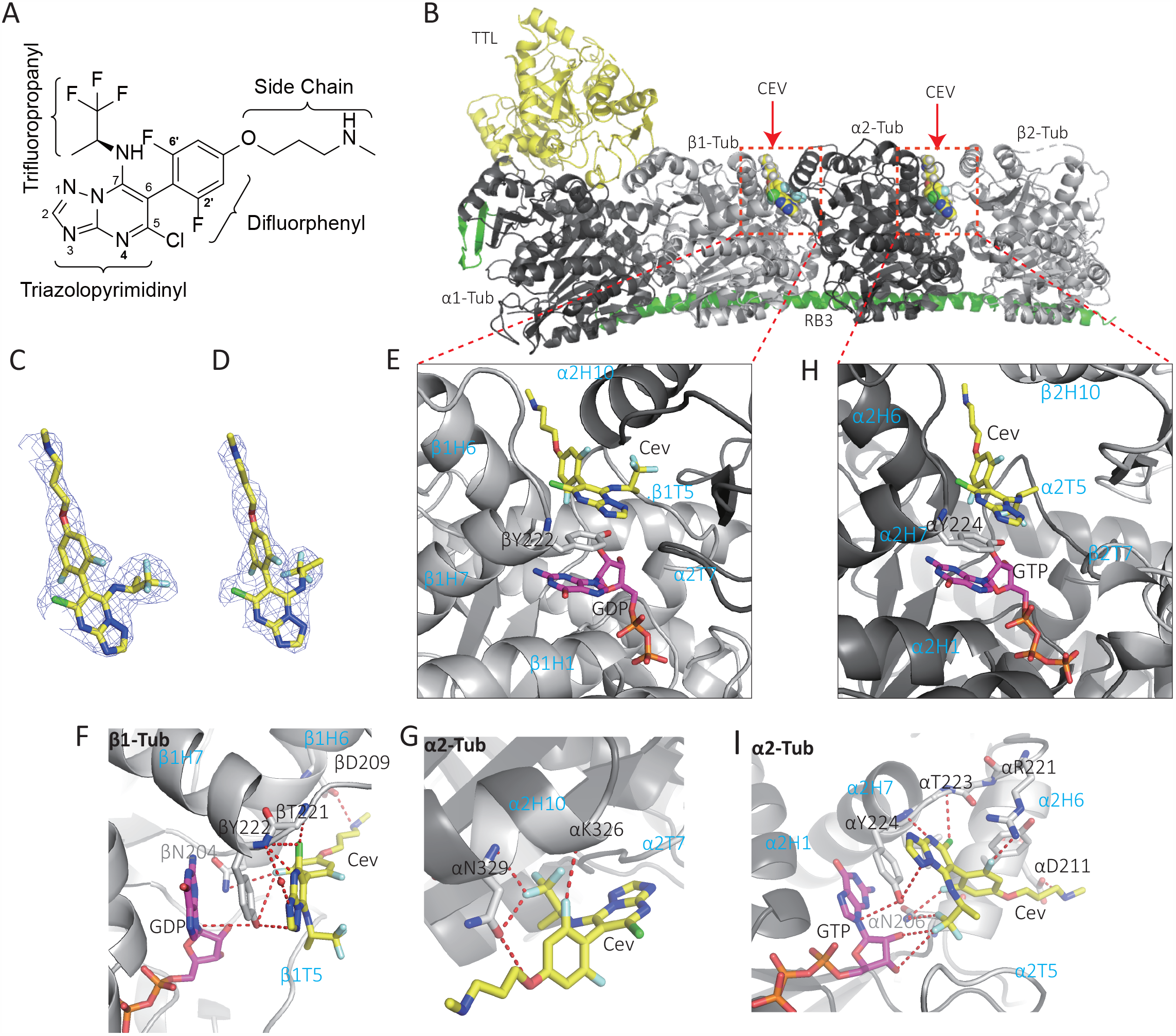
Crystal structure of cevipabulin-tubulin complex. (**A**) Chemical structure of cevipabulin. (**B**) Overall structure of cevipabulin-tubulin complex. TTL is colored yellow, RB3 is green, α-tubulin is black and β-tubulin is grey. Cevipabulin on β1-tubuin and α2-tubulin are all shown in spheres and colored yellow. (**C, D**) Electron densities of cevipabulins on (**C**) β1-tubulin or (**D**) α2-tubulin. The Fo-Fc omit map is colored light blue and contoured at 3d. (**E, F, G**) Close-up view of vinblastine-site cevipabulin binding to (**E, F**) β1-tubulin or (**G**) α2-tubulin. GDP or GTP is shown in magenta sticks. Cevipabulin is shown in yellow sticks. Side chain of β1-Y224 or α2-Y224 is show in grey sticks. (**H**) Interactions between the seventh-site cevipabulin and α2-tubulin. Coloring is the same as in (**E**). Residues from tubulin that form interactions with vinblastine-site cevipabulin are shown as sticks and labeled. Hydrogen bonds are drawn with red dashed lines. (**I**) Interactions between α2-tubulin and the seventh-site cevipabulin, color is the same as in (**F**), residues from tubulin that form interactions with the seventh-site cevipabulin are shown as sticks and labeled. Hydrogen bonds are drawn with red dashed lines. Cev: cevipabulin.

The binding region of cevipabulin in the vinblastine site was formed by residues from βH6, βH7, βT5 loop, αH10 and αT7 loop (Fig.2E). As presented in Figure 2F, the side chain of βY222 made π-π stacking interactions with triazolopyrimidinyl group of cevipabulin and the guanine nucleobase of GDP. Seven hydrogen bonds (N1 atom to side chain of βY222; N3 atom to main-chain nitrogen of βY222 through a water; N4 atom to main-chain nitrogen of βY222; 5-chlorine atom to both main-chain nitrogen of βY222 and βT221; 2’-fluorine atom to site chain of βY222 and main-chain nitrogen of βN204) between cevipabulin and β1-tubulin were observed. The -NH-group on the cevipabulin side chain formed a salt bridge with βD209. Besides, cevipabulin also exhibited four hydrogen bonds with α2-tubulin (oxygen atom on side chain to the side chain of αN329; 2’-fluorine atom to the main-chain nitrogen of αN326; one fluorine atom of trifluoropropanyl to both main and side chain of αN326) (Fig.2G).

The seventh site on α2-tubulin is formed by residues from αH1, αH6, αH7 and αT5 (Fig.2H). Similar to the vinblastine site, triazolopyrimidinyl of cevipabulin at this site also made π-π stacking interactions with the side chain of αY224 and the guanine nucleobase of GTP (Fig. 2I). There were eight hydrogen bonds (N1 atom to side chain of αY224; N4 atom to main-chain nitrogen of αY224; 5-chlorine atom to main-chain nitrogen of αT223; 2’-fluorine atom to site chain of αN206; 6’-fluorine atom to site chain of αR221; One fluorine atom of trifluoropropanyl to side chain of αN206;

Another fluorine atom of trifluoropropanyl to both O2’ and O3’ of GTP) between cevipabulin and α2-tubulin and a salt bridge between the -NH-group of cevipabulin side chain and αD211 (Fig. 2I). Notably, there is no hydrogen bond between cevipabulin and β2-tubulin at this new site.

### Direct biochemical assay confirms the binding of Cevipabulin to vinblastine site and the seventh site of tubulin

To ensure the new binding site is not an artefactual interaction due to high concentrated environment of the crystal, we further measured the binding of cevipabulin to tubulin heterodimer in solution (without the other proteins used to obtain crystals). Tubulin (20μM) in solution was incubated with different concentrations of cevipabulin and the content of bound cevipabulin were collected and quantified by liquid chromatography-tandem mass spectrometry (LC-MS/MS). As shown in Figure 3A, at respectively 64, 128 and 256 μM concentrations of cevipabulin, the stoichiometry was 1.74, 1.91 and 2.19 cevipabulin molecule per tubulin dimer, suggesting the formation of a 2:1 cevipabulin-tubulin complex. Since eribulin is a strong tubulin inhibitor binding to the microtubule plus ends (the vinblastine site) and can keep tubulin dimers in an assembly incompetent dimer state [4]. To confirm that cevipabulin simultaneously binds to the vinblastine site and the new binding site, we first blocked the vinblastine site with excessive eribulin before incubated with different concentrations of cevipabulin, and then the contents of both bound eribulin and cevipabulin were collected and quantified by LC-MS/MS. As shown in Figure 3B, at 24, 48 and 96 μM of cevipabulin, respectively, we observed that each tubulin dimer binds approximately one cevipabulin and one eribulin molecule, suggesting the formation of a 1:1:1 cevipabulin-eribulin-tubulin complex. Using the eribulin incubated tubulin, we could direct measure the dissociation constant (*K*_*d*_) of cevipabulin to the seventh site by a microscale thermophoresis assay (MST). As presented in Figure 3C, the MST results showed the *K*_*d*_ of cevipabulin to the seventh site is 0.97 ± 0.15 μM. We also solved the crystal structure of cevipabulin-eribulin-tubulin complex using the T2R-TTL crystal. As shown in Figure 3D, we observed one molecule of cevipabulin binding on β1-tubulin, one molecule of cevipabulin on α2-tubulin and one molecule of eribulin on β2-tubulin. Focusing on the α2β2-tubulin dimer, we obtained a 1:1:1 cevipabulin-eribulin-tubulin complex, which was consistent with the results of our biochemical experiments. Next, we performed competition experiments of cevipabulin to BODIPY-Vinblastine to measure the *K*_*d*_ value of cevipabulin to the vinblastine site, which is determined to be 0.90 ± 0.24μM (Fig. S1A and S1B). All these results confirm that cevipabulin could bind simultaneously to the seventh site at low micromole concentrations, in addition to the vincristine site.

**Figure 3.**
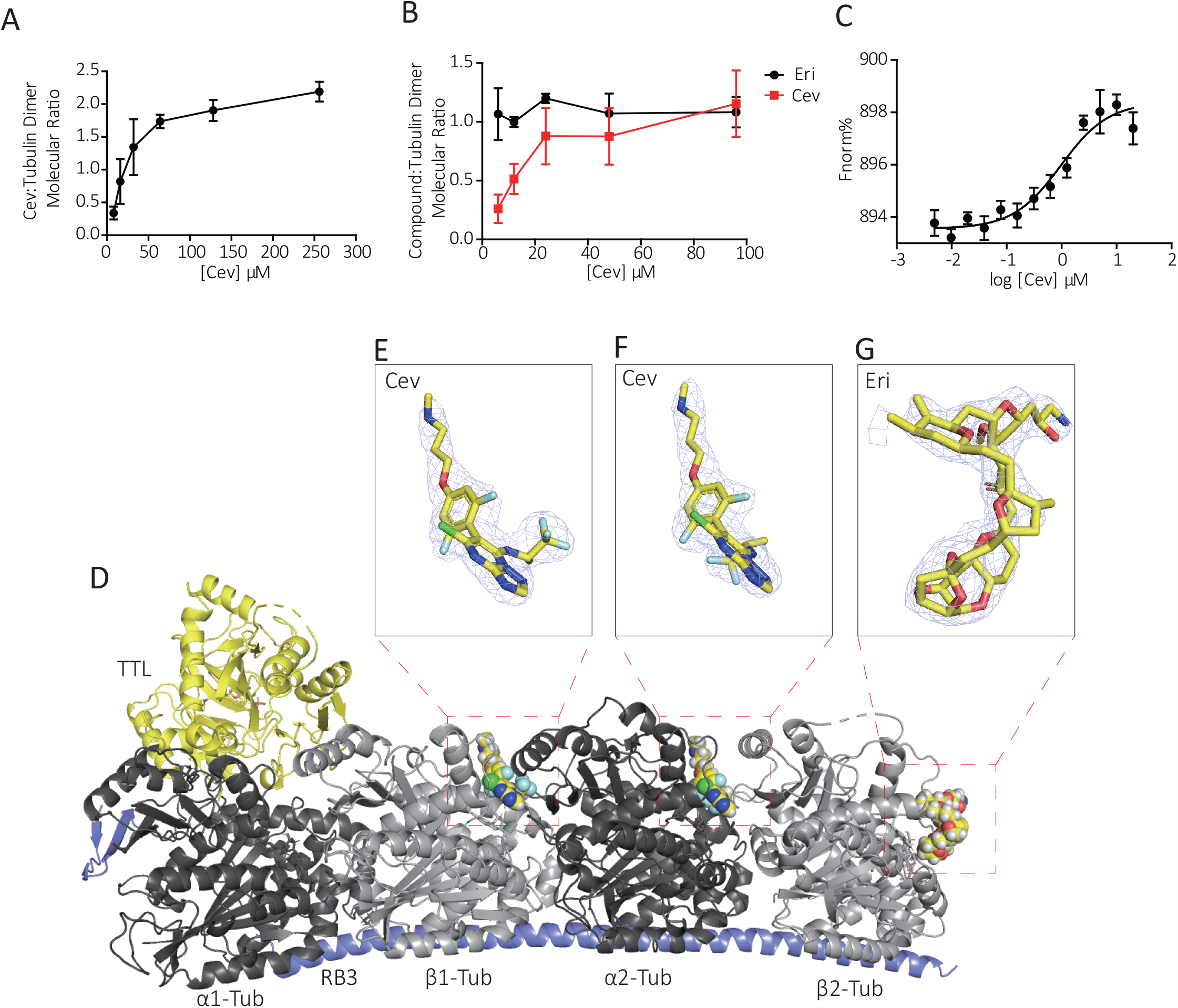
Measuring the binding stoichiometry of cevipabulin to tubulin. (**A**) Indicated concentrations of cevipabulin was incubated with tubulin (20μM) for 10min, and then the bound cevipabulin was quantified by LC-MS/MS. This graph presented the molecular ratio of cevipabulin: tubulin dimer. Data were shown as means ± SD of three independent experiments. (**B**) Indicated concentrations of cevipabulin was incubated with eribulin-tubulin complex (20μM) for 10min, and then the bound cevipabulin and eribulin was quantified by LC-MS/MS. This graph presented the molecular ratio of cevipabulin or eribulin: tubulin dimer. Data were shown as means ± SD of three independent experiments. (**C**) Binding of cevipabulin to the seventh site of tubulin was determined with the microscale thermophoresis assay. Data points represent means ± SD of three technical replicates each. (**D**) Structure of cevipabulin-eribulin-tubulin complex. TTL is colored yellow, RB3 is blue, α-tubulin is black and β-tubulin is grey. Cevipabulin on β1-tubuin and α2-tubulin and eribulin on β2-tubulin were all shown in spheres and colored yellow. (**E-G**) Electron densities of cevipabulin on β1-tubulin (**E**) or α2-tubulin (**F**) and eribulin on β2-tubulin (**G**). The Fo-Fc omit map is colored light blue and contoured at 3d. Cev: cevipabulin; Eri: eribulin; Tub: Tubulin.

### Cevipabulin binding to the seventh site to induce tubulin degradation

We then investigated whether the new binding site of cevipabulin to tubulin mediated tubulin degradation. When vinblastine site was occupied by eribulin or vinblastine, cevipabulin still retain the tubulin-degradation effect (Fig. 4A and S2A). Since the αY224 at the seventh site made π-π stacking interactions with triazolopyrimidinyl group of cevipabulin and the guanine nucleobase of GTP, it may be important for the binding of cevipabulin. Thus, single amino acid substitution (Y224G on α-tubulin) was employed to block the seventh site. When the seventh site was mutant, cevipabulin lost its ability to degrade tubulin (Fig.4B). These data indicated that the cevipabulin binding to the seventh site rather than vinblastine binding, plays a role in the degradation of tubulin. For further understand underline degradation mechanism, we synthesized two reported cevipabulin analogues (compounds **1** [16] and **2** [17]**) (**Fig. 4C). Compared with cevipabulin, compound **1** had no the N-substituted side chain and the trifluoropropanyl in compound **1** was replaced by an azabicyclo to obtain compound **2**. As shown in Fig.4D, compound **1** induced tubulin degradation, while compound 2 did not. LC/MS/MS results indicated that compound **1** bound to tubulin dimer with a 1:1 stoichiometry (Fig.4E). Further experiments indicated that αY224G mutation, but neither vinblastine nor compound **2**, inhibited compound **1** induced tubulin degradation (Fig. 4F, S2B and S2C). These results together demonstrated that cevipabulin or compound **1** binding to the seventh site induced microtubule degradation, while compound 2 only binding to vinblastine did not induce microtubule degradation, suggesting that trifluoropropyl of cevipabulin plays a key role at the new binding site.

**Figure 4.**
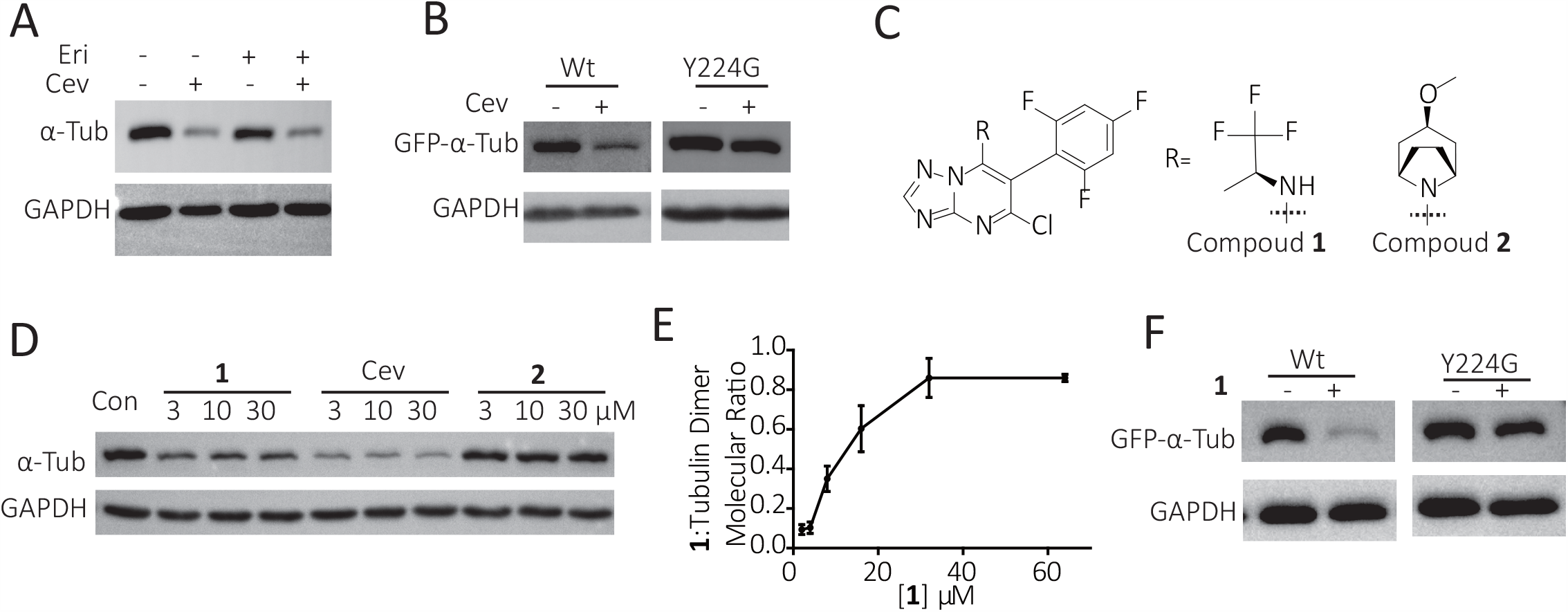
Cevipabulin binds to the seventh site to induce tubulin degradation. (**A**) HeLa cells were treated with 10 μM Eribulin for 1 h and then further treated with 1 μM cevipabulin for 16 h. The α-tubulin protein level was detected by immunoblotting. Results are representative of three independent experiments. (**B**) Vectors expressing either wild type or Y224G mutant GFP-tubulin were transfected to HeLa cells. After 24 hours, cells were treated with or without 1μM cevipabulin for 16 h. Then the protein level of GFP-α-tubulin was detected by immunoblotting. Results are representative of three independent experiments. (**C**) Chemical structure of cevipabulin derivatives. (**D**) Hela cells were treated with indicated compounds for 16 h. Then the protein level of α-tubulin was detected by immunoblotting. Results are representative of three independent experiments. (**E**) Indicated concentrations of compound **1** was incubated with tubulin (20μM) for 10min, and then the bound compound 1 was quantified by LC-MS/MS. This graph presented the molecular ratio of compound **1**:tubulin dimer. Data were shown as means ± SD of three independent experiments. (**F**) Vectors expressing either wild type or Y224G mutant GFP-tubulin were transfected to HeLa cells. After 24 hours, cells were treated with or without 10 μM compound **1** for 16 h. Then the protein level of GFP-α-tubulin was detected by immunoblotting. Results are representative of three independent experiments. Cev: cevipabulin. **1**: compound **1**; **2**: compound **2**; Vin: vinblastine.

### Cevipabulin and compound 1 destabilize tubulin by making the non-exchangeable GTP exchangeable

Crystal structures of cevipabulin-tubulin and compound **2**-tubulin (PDB code: 5NJH) could be superimposed very well in whole (Fig S3A, with a root-mean-square deviation (RMSD) of 0.45 Å over 1,930 Cα atoms) or in the vinblastine-site region (Fig S3B). Thus, the main conformational change of cevipabulin-tubulin is identical to compound **2**-tubulin, which has been described in detail in previous studies, except for the presence of additional density at the seventh site. We then focus on the study of this novel site. The non-exchangeable GTP plays a structural role and is important for the stability of tubulin dimers [19, 20]. As tubulin degradation induced by cevipabulin is mediated by its binding to the seventh site located near the non-exchangeable GTP, we suspect that cevipabulin and compound **1** may directly mediate the stability of tubulin. Using a thiol probe, tetraphenylethene maleimide (TPE-MI), which is non-fluorescent until conjugated to a thiol (Fig.5A) [21], we measured whether these compounds promote unfolding of tubulin. As shown in Figure 5B, TPE-MI alone did not increase fluorescence of tubulin while addition of 4M guanidine hydrochloride (non-selective protein denaturant) significantly increased fluorescence. Cevipabulin and compound **1** obviously increased tubulin fluorescence while vinblastine and compound **2** had no such effects, demonstrating that cevipabulin or compound **1** could promote unfolding of tubulin. In the seventh site, α-T5 loop undergoes a large outward shift after cevipabulin binding (Fig.5C), thus some important hydrogen bonds between the non-exchangeable GTP and α-T5 loop were destroyed (Fig. 5D), which we believe may affect the affinity of GTP with α-tubulin protein. We incubated tubulin with different compounds for 10 min at buffer containing 1mM GDP, and then the content of bound GTP and GDP were detected by LC-MS/MS. As presented in figure 5F, compared with the control groups, the GTP content became less while the GDP content increased after cevipabulin or compound **1** treated, in contrast, both the content of GTP and GDP had no change after compounds 2 or vinblastine treated, suggesting that cevipabulin or compound **1** binding reduces the affinity of this non-exchangeable GTP to α-tubulin and eventually makes it exchangeable. Therefore, the degradation process can be summarized as follows: cevipabulin or compound 1 binding at the seventh site push the α-T5 loop outward, breaking some important hydrogen bonds between α-T5 and the non-exchangeable GTP, lowering the affinity of this GTP to -Tubulin, thus reducing the stability of tubulin and leading to its destabilization and degradation.

**Figure 5.**
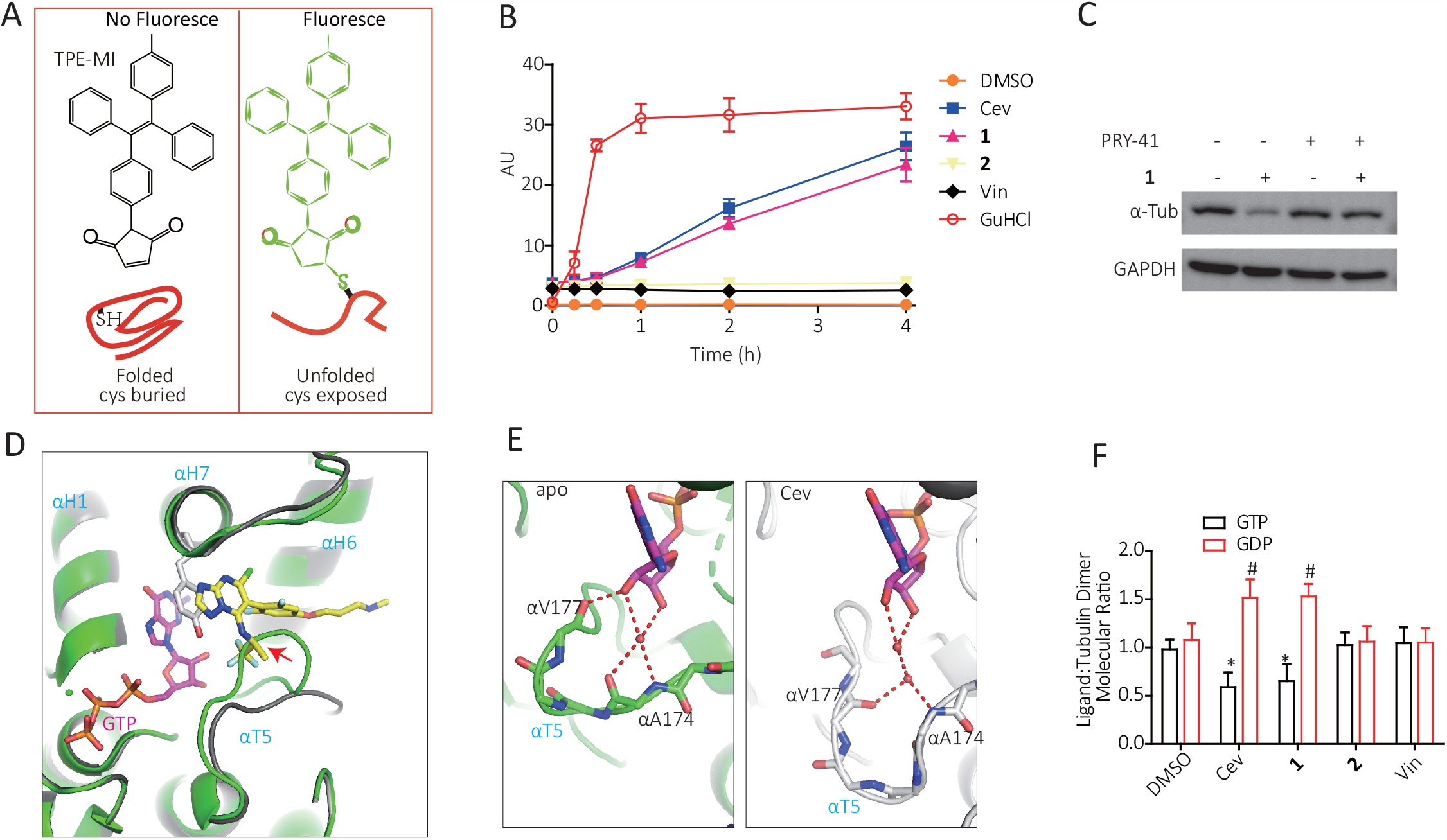
Cevipabulin or compound 1 decrease tubulin stability to promote tubulin destabilization and degradation. (**A**) The fluorescence of tetraphenylethene maleimide (TPE-MI) is enabled upon conjugation to a cysteine residue of a denatured protein. (**B**) Tubulin unfolding detected by TPE-MI. Tubulin (0.2 mg/ml) in PIPES buffer was mixed with 50 μM TPE-MI and the indicated compounds for various time before fluorescence (Excitation wavelength: 350nm; Emission wavelength: 470nm) were detected. Data were shown as means ± SD of three independent experiments. (**C**) Hela cells were treated with or without PYR-41(20 μM) for 1 hour before treated with 10 μM compound **1** for 16 h. Protein level of α-tubulin were detected by immunoblotting. Results are representative of three independent experiments. (**D**) Cevipabulin-tubulin (black) and apo-tubulin (green, PDB code:4i55) complexes were aligned on α2-tubulin and the close-up view of the seventh site were shown. GTP was shown in magenta sticks. Cevipabulin is shown in yellow sticks. Side chain of α2-Y224 is shown in grey sticks. The collision of α-T5 loop in apo-tubulin complex to cevipabulin is marked with Red arrow. (**E**) Left: Close-up view of the interaction between the non-exchangeable GTP and the α-T5 loop in apo-tubulin complex. GTP was shown in magenta sticks. The mainchain of α-T5 loop is shown in sticks. Hydrogen bonds are drawn with red dashed lines. Right: Close-up view of the interaction between the non-exchangeable GTP and the α-T5 loop in cevipabulin-tubulin complex. GTP was shown in magenta sticks. The mainchain of α-T5 loop is shown in sticks. Hydrogen bonds are drawn with red dashed lines. (**F**) Tubulin (20 μM) in PIPES buffer supplemented with 1 mM GDP were incubated with cevipabulin (30μM), compound 1 (30μM), compound 2 (30μM) or vinblastine (30μM) for 10min, the bound GTP and GDP were further quantified with an LC-MS/MS method. This graph presented the molecular ratio of GTP or GDP: tubulin dimer. Data were shown as means ± SD of three independent experiments. *p<0.05, GTP content as compared to the DMSO treated group; #p<0.05, GDP content as compared to the DMSO treated group. Cev: cevipabulin; Vin: vinblastine. **1**: compound **1;2**: compound **2**.

## Discussion

Our study identifies a novel binding site on α-tubulin, the seventh site. As this new site is located near the non-exchangeable GTP site which is important for tubulin stability [19, 20, 22], inhibitors such as cevipabulin and compound **1** binding to the seventh site reduce tubulin stability and promote tubulin degradation. This novel site on α-tubulin is spatially corresponding to the vinblastine site on β-tubulin, which is also bound by cevipabulin. The binding pocket of cevipabulin to these two sites is very similar (formed by αH1, αH6, αH7, αT5 for the seventh site and βH1, βH6, βH7, βT5, αH10, αT7 for the vinblastine site) and the binding modes of cevipabulin are also similar except the trifluoropropanyl of cevipabulin adopts different conformations. Vinblastine-site cevipabulin is mainly located on β1-tubulin and makes lots of hydrogen bonds with β1-tubulin while its trifluoropropanyl is oriented towards α2-tubulin and makes four hydrogen bonds interactions with α2-tubulin. The-seventh-site cevipabulin is totally located on α2-tubulin and makes lots of hydrogen bond with α2-tubulin and its trifluoropropanyl is also oriented towards α2-tubulin to establish hydrogen bonds with the non-exchangeable GTP. Of note, compound **2** lacking the trifluoropropanyl could not bind to the seventh site and showed no tubulin degradation effect, suggesting the trifluoropropanyl-GTP interaction is important for cevipabulin binding to the seventh site. We noticed that in the compound **2**-tubulin complex, although compound **2** bound only to the vinblastine site, the αT5 loop at the seventh site also had an outward shift as in the cevipabulin-tubulin complex. It seems to suggest that that compound 2 could also bind to the seventh site but with a low affinity and density that is undetectable. However, biochemical results indicated that compound **2** showed no interaction with the seventh site at low micromole concentrations [17] and compound **2** cannot induce tubulin degradation. Although we confirmed that compound **1** binds only to the seventh site and not to the vinblastine site, we unfortunately did not obtain the crystal structure of tubulin-compound **1** complex (possible due to the lower affinity of compound **1** to the seventh site), which might provide other vital information of the seventh site.

To prove the binding of cevipabulin to the seventh site is not an artefactual interaction due to the high concentration of the crystal environment or as a result of crystal packing or an artefact of T2R-TTL protein complex used, we measured the binding in solution (without the RB3 and TTL protein used for crystallization) at micromole concentration. We observed that 20 μM tubulin dimer in solution could be bound with 40μM cevipabulin at most, suggesting the formation of a 2:1 cevipabulin-tubulin complex. In addition, when the vinblastine site was blocked by a strong tubulin inhibitor-eribulin, 20 μM tubulin dimer can still be bound with 20 μM cevipabulin and forms a 1:1:1 cevipabulin-eribulin-tubulin complex. All these demonstrated that cevipabulin can bind to the seventh site at low micromole concentration and an MST assay directly determined the dissociation constant (*K*_*d*_) of cevipbulin to the seventh site (eribulin-tubulin complex) to be 0.97±0.15 μM. We further soaked both cevipabulin and eribulin simultaneously to T2R-TTL complex to obtain a cevipabulin-eribulin-tubulin complex. We observed one molecule cevipabulin on α2-tublin, and one molecule eribulin on β2-tububulin (Figure 2D), indicating the formation of a 1:1:1 cevipabulin-eribulin-tubulin complex that is in consistent with the biochemical results. Unexpectedly, we found one molecule cevipabulin but not eribulin on β1-tublin, suggesting cevipabulin has much higher affinity to the interdimer-face vinblastine site than eribulin. Thus, though these two compounds both bind the vinblastine site, eribulin preferred binds to the vinblastine site at the plus end while cevipabulin only binds to interdimer-face vinblastine site with high affinity.

The novel site is located near the non-exchangeable GTP, which plays a structural role and is important for the stability of tubulin dimers [19, 20]. This non-exchangeable GTP forms a number of hydrogen bonds with surrounding amino acid residues and a magnesian ion [19]. Single mutation abolishing hydrogen bond with this GTP could reduce the affinity of GTP and the absence of the magnesian ion would reduce the protein stability [19, 22]. Cevipabulin binding to the novel site pushes α-T5 loop outward and breaks several key hydrogen bonds, reduces the non-exchangeable to α-tubulin and promote its unfolding, then inducing its degradation in a proteasome-dependent pathway. Therefore, compounds binding the seventh site may all possess tubulin degradation effect because the α-T5 loop outward shift was necessary for compounds binding.

Here, we reported a novel binding site on α-tubulin that possessed tubulin degradation effect that was distinct from the traditional MDAs and MSAs. Using this specific site, a new class of tubulin degraders can be designed as anticancer drug targeting α-tubulin.

## Materials and Methods

### Reagents

Colchicine, Vinblastine, BODIPY-vinblastine, β,γ-Methyleneadenosine 5′-triphosphate disodium salt (AMPPCP), Tetraphenylethene maleimide (TPE-MI), and DL-dithiothreitol (DTT) were purchased from Sigma; Eribulin mesylate was obtained from MOLNOVA. Guanidine hydrochloride, MG132 and PYR-41 were obtained from Selleck; Cevipabulin was from MedChemExpress; Purified tubulin was bought from Cytoskeleton, Inc.; Antibodies (α-tubulin antibody, β-tubulin antibody, GAPDH antibody and gout anti mouse second antibody) were bought from Abcam.

### Chemistry

All the chemical solvents and reagents used in this study were analytically pure without further purification and commercially available. TLC was performed on 0.20 mm silica gel 60 F_254_ plates (Qingdao Ocean Chemical Factory, Shandong, China). Visualization of spots on TLC plates was done by UV light. NMR data were measured for ^1^H at 400 MHz on a Bruker Avance 400 spectrometer (Bruker Company, Germany) using tetramethylsilane (TMS) as an internal standard. Chemical shifts were quoted in parts per million. High Resolution Mass Spectra (HRMS) were recorded on a Q-TOF Bruker Daltonics model IMPACT II mass spectrometer (Micromass, Manchester, UK) in a positive mode.

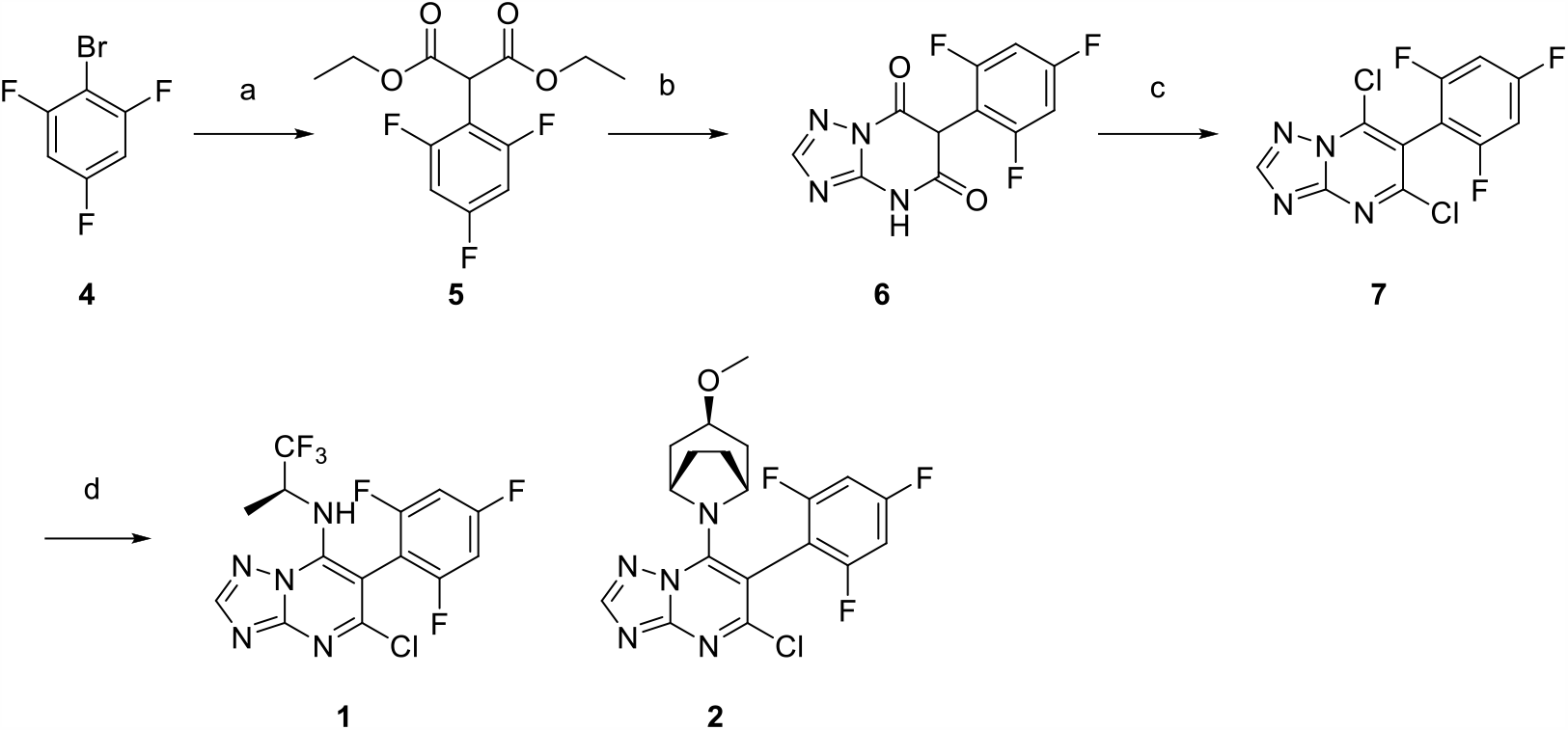

*Scheme1: Reagents and conditions: a) diethyl malonate, NaH, CuI, dioxane, r*.*t*.*-reflux; b) 3-amino-1,2,4-triazole, tributylamine, 180 °C; c) POCl*_*3*_, *reflux; d) amine, K*_*2*_*CO*_*3*_, *DMF, r*.*t*.

General procedure for the synthesis of diethyl 2-(2,4,6-trifluorophenyl)malonate(**5**)

To a stirred solution of diethyl malonate (320 mg, 2.0 mmol) in 1,4-dioxane was added 60% sodium hydride (96 mg, 2.4 mmol) by portions at room temperature. Then cupper (I) bromide (380 mg, 2.0 mmol) and compound **4** (211mg, 1.0 mmol) was added. The reaction mixture was stirred at room temperature for 30 minutes and then refluxed for 8 hours under nitrogen protection. After completion of the reaction, the mixture was cooled to room temperature and hydrochloric acid (12 N, 50 mL) was added slowly. The organic phase was separated off and the aqueous phase was extracted with ethyl acetate (×2). The combined organic phase was concentrated in vacuo. The residue was purified by chromatograph on silica gel with petroleum ether and ethyl acetate as eluent to give compound **5** as a white solid. Yield: 62%. ^1^H NMR (400 MHz, DMSO) δ 7.36 – 7.18 (m, 2H), 5.15 (s, 1H), 4.18 (q, *J* = 7.1 Hz, 4H), 1.23 – 1.14 (m, 6H). HRMS-ESI: calcd for [C_13_H_13_F_3_O_4_+Na]^+^ 313.0664, found: 313.0663.

General procedure for the synthesis of 5,7-dichloro-6-(2,4,6-trifluorophenyl)-[1,2,4]triazolo[1,5-*a*]pyrimidine (**7**)

A mixture of 3-amino-1,2,4-triazole (84 mg, 1.0 mmol), compound **5** (290 mg, 1.0 mmol) and tributylamine (1.0 mL) was heated at 180 °C for 4 hours. After the reaction mixture was cooled to room temperature, the residue was diluted with dichloromethane, washed with dilute hydrochloric acid and water and crystallized from diisopropyl ether to yield 116 mg of compound **6** (brown solid, 41% yield). Then phosphorus oxitrichloride (10 mL) was added to a 25 mL round-bottom flask filled with compound **6** (282 mg, 1.0 mmol), and refluxed for 4 hours. After completion of the reaction, the reaction mixture was cooled to room temperature and the solvent was distilled off. The residue was diluted with water and ether acetate. The organic phase was separated, washed with dilute sodium bicarbonate solution and brine, dried, concentrated in vacuo and purified by chromatograph on silica gel with petroleum ether and ethyl acetate as eluent to give compound **7** as a white solid. Yield: 66%, ^1^H NMR (400 MHz, DMSO) δ 8.90 (s, 1H), 7.62-7.55 (m, 2H). HRMS-ESI: calcd for [C_11_H_3_Cl_2_F_3_N+H]^+^ 318.9765, 320.9736, found: 318.9764, 320.9739; calcd for [C_11_H_3_Cl_2_F_3_N+Na]^+^ 340.9585, 342.9555, found: 340.9576, 342.9565.

General procedure for the synthesis of **1-2** Compounds **1** and **2** were prepared as described in Zhang et al[23]. Compound **7** (160 mg, 0.5 mmol), (*S*)-1,1,1-trifluoropropan-2-amine hydrochloride (75 mg, 0.5 mmol, for **1**), or (1R,3r,5S)-3-methoxy-8-azabicyclo[3.2.1]octane (71 mg, 0.5 mmol for **2**), and potassium carbonate (276mg, 2.0 mmol) was dissolved in DMF (5 mL) and stirred at room temperature for 4 hours. After completion of the reaction, water and ethyl acetate was added. The organic phase was separated, washed with brine, dried over anhydrous sodium sulfate, concentrated in vacuo and purified by chromatograph on silica gel with petroleum ether and ethyl acetate as eluent to give compounds **1 and 2** as white solid. Yield: 48%-63%.

(S)-5-chloro-6-(2,4,6-trifluorophenyl)-N-(1,1,1-trifluoropropan-2-yl)-[1,2,4]triazolo[1,5-a]pyrimidin-7-amine (**1**)

Yield: 48%, ^1^H NMR (400 MHz, CDCl_3_) δ 8.40 (s, 1H), 6.93-6.89 (m, 2H), 5.96 (d, *J* = 10.6 Hz, 1H), 4.75 (s, 1H), 1.43 (t, *J* = 10.0 Hz, 3H). HRMS-ESI: calcd for [C_14_H_8_ClF_6_N_5_+H]^+^ 396.0451, found 396.0488; calcd for [C_14_H_8_ClF_6_N_5_+Na]^+^ 418.0270, found 418.0263.

5-chloro-7-((1*R*,3*r*,5*S*)-3-methoxy-8-azabicyclo[3.2.1]octan-8-yl)-6-(2,4,6-trifluorophenyl)-[1,2,4]triazolo[1,5-*a*]pyrimidine (**2**)

Yield: 63%, ^1^H NMR (400 MHz, DMSO) δ 8.57 (s, 1H), 7.52-7.48 (m, 2H), 4.58 (s, 2H), 3.43 (t, *J* = 4.0 Hz, 1H), 3.17 (s, 3H), 2.01 (dt, *J* = 10.2, 5.1 Hz, 4H), 1.90 (d, *J*= 14.6 Hz, 2H), 1.77 – 1.67 (m, 2H). HRMS-ESI: calcd for [C_19_H_17_ClF_3_N_5_O+H]^+^ 424.1152, found 424.1152; calcd for [C_19_H_17_ClF_3_N_5_O+Na]^+^ 446.0971, found 446.0964.

### Cell culture

HeLa, Hct116, H460 and SU-DHL-6 cells were all sourced from American Type Culture Collection. H460 cells were cultured in RPMI 1640 medium and HeLa, Hct116 and SU-DHL-6 cells were cultured in Dulbecco’s Modified Eagle’s medium. Both media were supplemented with 5%-10% fetal bovine serum and about 1% penicillin-streptomycin. The culture temperature was set at 37°C, and cells were grown in a humidified incubator with 5% CO_2_. All cells have been authenticated by STR tests and are free of mycoplasma.

### Label free Quantitative Proteomics

HeLa cells were treated with or without 1μM cevipabulin for six hours and then all cells were collected and lysed with radioimmunoprecipitation assay buffer (containing proteinase inhibitor mixture) for 30min on ice. Then all samples were centrifuged at 10,000 g for 30 minutes to pellet cell debris. Supernatants were collected and stored at −80°C before analysis. We have done three biological repeats. Then the following label-free quantitative proteomic analysis of these samples were carried out following the procedure as described previously[13].

### Immunoblotting

Cells were plated on six-well plates and cultured for 24 hours before treated with different compounds for different time. Total cells were harvested and washed by phosphate buffer saline (PBS) before centrifuged at 1000 g for 3min. Then 1╳loading buffer (diluted from 6╳loading buffer by radioimmunoprecipitation assay buffer l) was added to the cell pellets and lysed for 10min. Samples were then incubated in boiling water for 10 min and then stored at −20°C before use. Equal volume of samples was loaded to 10% SDS-PAGE for electrophoresis and then transferred to a polyvinylidene difluoride (PVDF) membranes at 4°C for 2 hours. Proteins on PVDF membranes were incubated in blocking buffer (5% skim milk diluted in 1╳PBST(PBS buffer with 0.1% Tween-20)) for 1hours. Then the PVDF membranes were incubated with first antibodies (dilute ed in blocking buffer) for 12hours and washed for three times with PBST before incubated with second antibody (diluted in blocking buffer) for 45 min and washed for three times with PBST again. At last, the PVDF membranes were immersed in enhanced chemiluminescence reagents for 30 seconds subjected to image with a chemiluminescence image analysis system (Tianneng, China).

### Quantitative-PCR

HeLa and Hct116 cells were plated on six-well plates and culture for 24hours before treated with cevipabulin for different time. Total mRNA of both HeLa and Hct116 cells were extracted with TRIzol (Invitrogen, USA) agents following the manufacturer’s protocol and then qualified using a NanoDrop1000 spectrophotometer (Thermo Fisher Scientific, USA. The cDNA synthesis was carried out using a high Capacity cDNA Reverse Transcription Kit (Applied Biosystems, USA). Taq Universal SYBR Green Supermix (BIO-RAD, USA) was employed for further Quantitative PCR analysis on a CFX96 Real-time PCR System (BIO-RAD, USA). Relative mRNA level of both *α-tubulin* and *β-tubulin* were normalized to that of GAPDH. Theprimers employed were as follows:

*α-tubulin*: forward primer, TCGATATTGAGCGTCCAACCT; reverse primer, CAAAGGCACGTTTGGCATACA;

*β-tubulin*: forward primer, TGGACTCTGTTCGCTCAGGT; reverse primer, TGCCTCCTTCCGTACCACAT;

*GAPDH*: forward primer, GGAGCGAGATCCCTCCAAAAT; reverse primer, GGCTGTTGTCATACTTCTCATGG.

### Single amino acid substitution on α-tubulin

The pIRESneo-EGFP-alpha Tubulin plasmid was obtained from Addgene (USA) and mutation (Y224G) of α-Tubulin were performed using a Q5 Site-Directed Mutagenesis kit (NEB #E0554S, USA). Hela cells were plated on six-well plates and incubated for 24 hours before transfected with these plasmids by Lipofectamine 2000. Then cells were culture for another 24 hours before treated with or without different compounds for 16 hours. Total protein was extracted and analyzed by immunoblotting to detect the content of GTP-α-tubulin and GAPDH was employed as loading control.

### Stoichiometry of cevipabulin or compound 1 to tubulin dimer

Tubulin (20μM) in PIPES buffer (80 mM PIPES pH 6.9, 0.5 mM EGTA, 2 mM MgCl_2_) supplemented with 1 mM GTP were incubated with cevipabulin (8, 16, 32, 64, 128 or 256 μM) or compound **1** (2, 4, 8, 16, 32 or 64 μM) for 10 min. Then the unbounded compounds were washed away with an ultrafiltration method (Centrifugation in a10 kd ultrafiltration tube at 13,000 rpm for 5 min for three times). The retentate (tubulin) was heated to 90 °C for 5min to denature the tubulin protein and release the bound cevipabulin. The bound cevipabulin was quantified with an LC-MS/MS.

### Stoichiometry of cevipabulin to eribulin preincubated tubulin dimer

Tubulin (1μM) in PIPES buffer (80 mM PIPES pH 6.9, 0.5 mM EGTA, 2 mM MgCl_2_) supplemented with 1 mM GTP were incubated with 1.5μM eribulin for 10min and then concentrated to 20 μM before further incubated with 6, 12, 24, 48 or 96 μM cevipabulin for another 10min. Then the unbounded compounds were washed with an ultrafiltration method (centrifugation in a10 kd ultrafiltration tube at 13,000 rpm for 5 min for three times). The retentate (tubulin) was heated to 90 °C for 5min to denature the tubulin protein and release the bound compounds. The bound cevipabulin and eribulin was quantified with an LC-MS/MS.

### LC-MS/MS method

LC-MS/MS analysis was performed on an ultrafast liquid chromatography system (Shimadzu) coupled with an AB SCIEX Qtrap 5500 mass spectrometer, equipped with electrospray ionization (ESI) source. Instruments control and collection of chromatographic and mass spectrometry information were carried out by analyst 1.6.2 software (AB SCIEX, USA). Chromatographic separation was achieved on a Waters ACQUITY UPLC BEH C_18_ column (2.1 mm × 100 mm I.D., 1.7 µm). The mobile phase system consisted of 0.1% formic acid in water (A) and Acetonitrile (B) using a gradient elution as follows: 0-1.0 min, 10-90% B; held 90% B for 1 min. The flow rate was 0.5 mL/min, and the temperature of the column and autosampler were maintained at 35 and 15 °C, respectively. The injection volume was 1µL. In the MS analysis, positive ionization mode was used for samples detection, with the following optimized mass spectrometric parameters: Ionspray voltage, 5500 V; Declustering Potential, 100 V; Temperature, 500 °C. Eribulin, compound **1**, cevipabulin and internal standard (IS) were conducted in multi reaction monitoring (MRM) mode with ions pair of 730.4>680.4, 396.0>360.1, 465.0>358.2 and 265.2>232.2, respectively.

### MST assay

Binding of cevipabulin to the seventh site was detected with an MST assay with a Monolith NT.115 instrument (NanoTemper Technologies). Purified tubulin was labeled using the Monolith protein labeling kit RED-NHS (NanoTemper Technologies) and then diluted to 200 nM concentration before incubated with 2 μM eribulin for 20min at room temperature. Different concentrations (20 μM to 48.9 nM) of cevipabulin were incubated with labeled tubulin (100 nM) in assay buffer (80 mM PIPES, pH 6.9, 0.5 mM EGTA, 2 mM MgCl_2_,1 mM GTP) for 10 min at 4 °C. Samples were loaded into glass capillaries for detection. *Kd* values were obtained using NanoTemper software.

### Determination of the dissociation constant of cevipabulin to vinblastine site by competition assay

Different concentrations of BODIPY-Vinblastine (50μM to 48.9nM) were incubated with 50 nM tubulin dimer and RB3 (T2R) complex (2: 1.2) plus with 1% DMSO or 50 μM vinblastine (final DMSO concentration was 1%) for 10min. The fluorescent emission ratio (520 nm/490 nm, 340 nm excitation) were detected using a Biotech Gen5 spectrophotometer (Biotek, USA). The data of each concentration were calculated as Data_DMSO_-Data_Vinblastine._ The specific equilibrium binding constant (*Kd*)was obtained by fitting the data into a one-site specific binding equation using Graphpad Prism 5.0. The *Kd* value was determined to be 1.571 ± 0.23 μM. T2R complex (50nM) was preincubated with 1μM BODIPY-Vinblastine for 10min before furture incubated with serious concentrations of cevipabulin for another 10min. Then fluorescent emission ratio (520 nm/490 nm, 340 nm excitation) were collected. The data were fit into a one-site competitive binding equation in a dose-dependent manner to get an IC_50_ values. The inhibition constant (Ki) value was calculated using the following equation [24]:

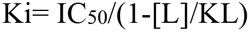

IC_50_ is the concentration of cevipabulin that inhibits 50% of bindin; [L] is the concentration of BODIPY-Vinblastine (1000 nM); KL is the Kd value of BODIPY-Vinblastine to T2R (1571 nM).

### Structural Biology

Protein expression and purification were detailly described in our precious study [25]. Tubulin, RB3 and TTL (2:1.3:1.2 molar ratio) were mixed together, then 5 mM tyrosine, 10 mM DTT and 1 mM AMPPCP were added and then the mixture was concentrated to about 15 mg/ml at 4 °C. The crystallization is conducted using a sitting-drop vapor-diffusion method under 20°C and the crystallization buffer is optimized as: 6% PEG4000, 8% glycerol, 0.1 M MES (pH 6.7), 30 mM CaCl_2_, and 30 mM MgCl_2_. Seeding method was also used to obtain single crystals. Crystals appeared in about 2-days and in a rod like shape and the size reached maximum dimensions within one week. About 0.1 μL cevipabulin (diluted in DMSO with a concentration of 100 mM) was added to a drop containing tubulin crystal and incubated for 16 h at 20 °C to get cevipabulin-tubulin complex. About 0.1 μL cevipabulin (100 mM) and 0.1 μL eribulin (100 mM) were added to a drop containing tubulin crystal and incubated for 16 h at 20 °C to get cevipabulin-eribulin-tubulin complex. The following data collection and structure determination were the same as previous description [25].

### TPE-MI as a thiol probe to detect unfolded protein

TPE-MI is a small molecule which is inherently non-fluorescent until covalently binds to a thiol by its maleimide [21, 26]. This molecule could be used to monitor purified protein unfolding *in vitro* [21]. Purified tubulin (0.2mg/ml) was diluted in PEM buffer supplemented with 1 mM GTP and then mixed with 50 μM TPE-MI and different compounds for various time. Then the samples were immediately subjected to a microplate reader (Biotek, USA) to detect the fluorescence (Excitation wavelength:350nm; Emission wavelength: 470nm).

### Quantification of tubulin bound GTP and GDP

Tubulin (20 μM) in PIPES buffer supplemented with 1 mM GDP were incubated with cevipabulin (30μM), compound **1** (30μM), compound **2** (30μM) or vinblastine (30μM) for 10min, and then tubulin was washed with PIPES buffer for three times in a 10-kd ultrafiltration tube. The retentate (tubulin) was heated to 90 °C for 5min to denature the tubulin protein and release the bound GTP and GDP which was further quantified with an LC-MS/MS method. Tubulin (20 μM) in PIPES buffer supplemented with 1 mM GDP were incubated with cevipabulin (30μM), compound **1** (30μM), compound **2** (30μM) or vinblastine (30μM) for 10min, and then tubulin was washed with PIPES buffer for three times in a 10-kd ultrafiltration tube. The retentate (tubulin) was heated to 90 °C for 5min to denature the tubulin protein and release th e bound GTP and GDP which was further quantified with an LC-MS/MS method.

### Statistical analysis

Data are presented as means. Statistical differences were determined using an unpaired Student’s t test. p values are indicated in figure legend when necessary: ^*^ or #, p< 0.05.

## Acknowledgement

This work received funds from National Natural Science Foundation of China (81872900, 81803021); 1.3.5 project for disciplines of excellence, West China Hospital, Sichuan University; National Major Scientific and Technological Special Project for ‘Significant New Drugs Development’ (2018ZX09201018-021); Post-Doctor Research Project, Sichuan University (2020SCU12036); China Postdoctoral Science Foundation Grants (2019T120855 and 2019M650248); Sichuan Science and Technology Program Grants (2019YJ0088, 2019YFH0123 and 2019YFH0124). Post-doctoral Research Project, West China Hospital, Sichuan University Grant 2018HXBH027.

## Author contributions

J.Y. performed most of the cellular and biochemical experiments and wrote the draft. J.Y., Y.Y., W.Y, L.N. and Q.C. performed the structural biology experiments. Y.L. synthesized all chemical compounds. H.Y., Y.Z., Z.W., Z.Y., H.P., H.W., M.Z., J. W. L.Y., and L.O., performed some of these biochemical experiments. M.T. performed the LC-MS/MS experiment. J.Y., W.L. and L.C. conceived the idea and supervised the study. J.Y., Q.C., W.L. and L.C. revised the manuscript.

All authors approved the final manuscript.

## Data availability

Atomic coordinates and structure factors of cevipabulin-tubulin and cevipabulin-eribulin-tubulin complexes have been deposited in the Protein Data Bank under accession codes 7CLD and 7DP8 respectively. Further information and requests for resources and reagents should be directed to and will be fulfilled by Jianhong Yang.

---

**Figure S1:**
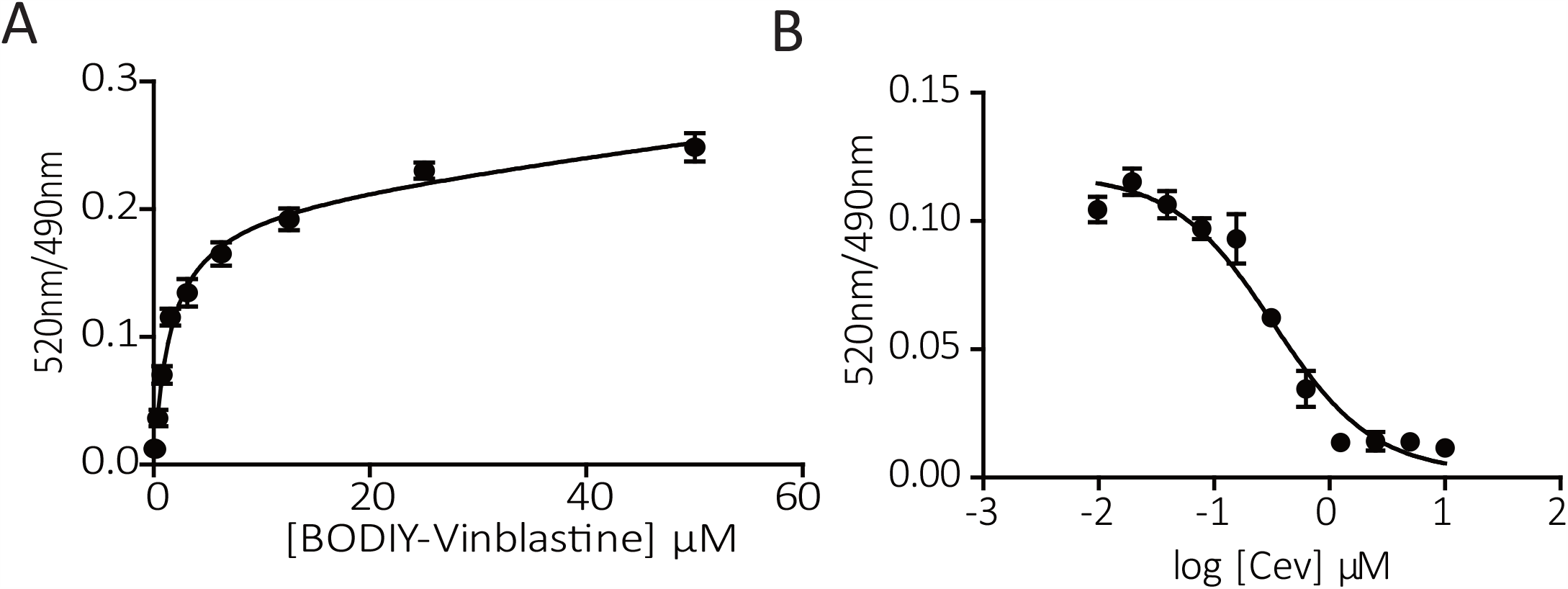
Determination of the equilibrium binding constant (*Kd*) of cevipabulin to the vinblastine site. **(A)** Binding curves of the indicated concentrations of BODIPY-Vinblastine to 50 nM T2R complex. The specific binding curve was generated by subtracting non-specific binding from total binding. The specific equilibrium binding constant (Kd) was calculated as 1.571 ± 0.23 μM. (**B**) The cevipabulin dose-response curves in the presence of 1 μM BODIPY-Vinblastine and 50 nM T2R complex. The data were fit into a one-site competitive binding equation in a dose-dependent manner to get an IC_50_ value, which was 0.3256 ±0.086 μM. Data points represent means ± SD of three technical replicates each. The inhibition constant (*Ki*) value was calculated using the following equation: Ki= IC_50_/(1-[L]/KL). Where IC_50_ is the concentration of cevipabulin that inhibits 50% of binding (0.3256 ±0.086 μM); [L] is the concentration of BODIPY-Vinblastine (1.0 μM); KL is the Kd value of BODIPY-Vinblastine to T2R (1.571 ± 0.23 μM). And the *Ki* value is calculated as 0.896 ± 0.24μM, which could also be regarded as the equilibrium binding constant (*Kd*).

**Figure S2.**
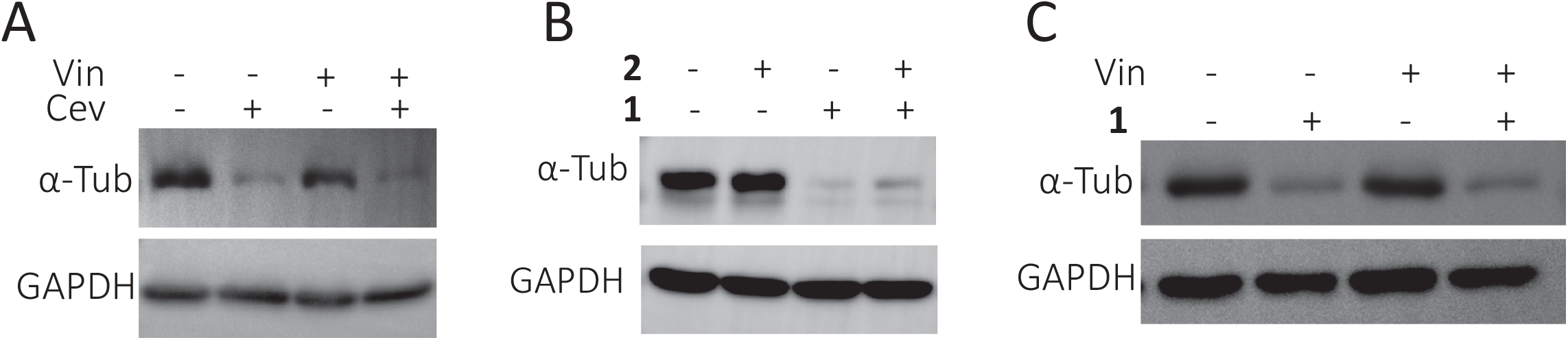
Compound 1 binds to the seventh site to induce tubulin degradation. (**A**) HeLa cells were treated with or without 10 μM vinblastine for 1 hour before treated with 1 μM cevipabulin for 16 h and then the protein level of α-tubulin was detected by immunoblotting. Results are representative of three independent experiments. (**B**) HeLa cells were treated with or without 30μM compound **2** for 1hour before treated with 10μM compound **1** for 16 h and then the protein level of α-tubulin was detected by immunoblotting. Results are representative of three independent experiments. (C) HeLa cells were treated with or without 10μM vinblastine for 1hour before treated with 10μM compound **1** for 16 h and then the protein level of α-tubulin was detected by immunoblotting. Results are representative of three independent experiments. Cev:cevipabulin; Vin:vinblastine; **1**: compound **1; 2**: compound **2**.

**Figure S3.**
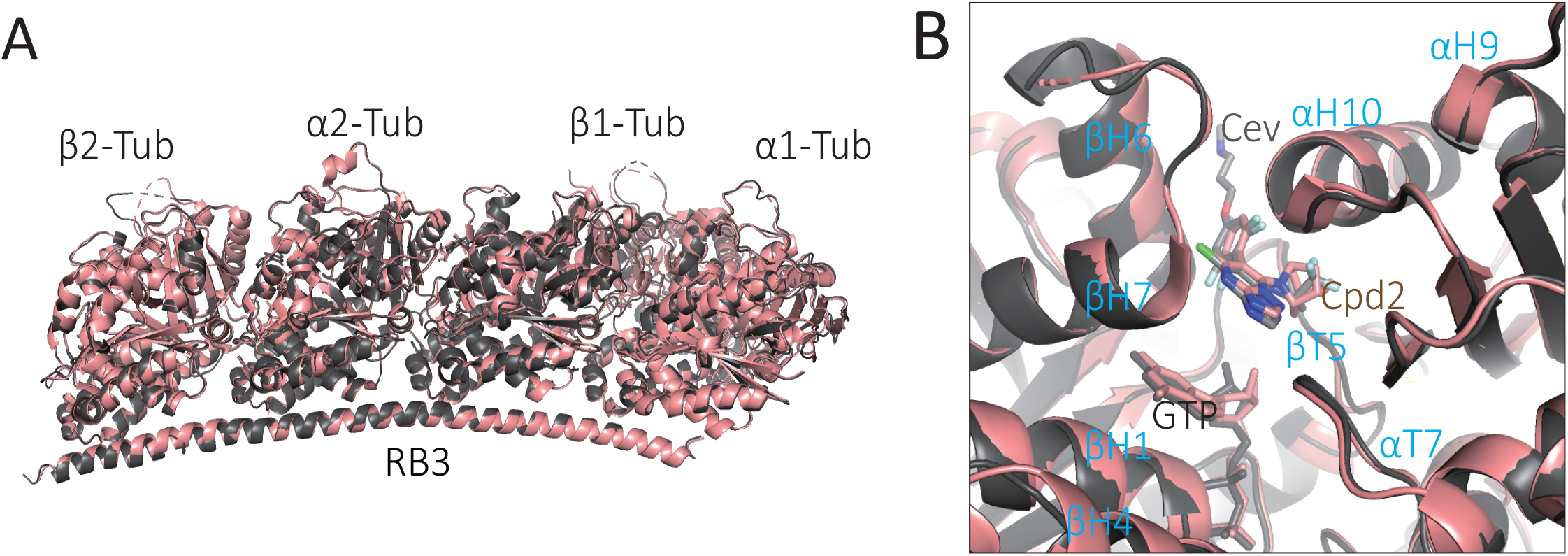
Aligned structure of cevipabulin-tubulin (black) and compound 2-tubulin (salmon) complexes. (**A**) Overview of the aligned structures of cevipabulin-tubulin complex (dark) and compound **2**-tubulin complex (salmon) (PDB code: 5NJH). (**B**) Close-up view of the cevipabulin and compound **2** binding to inter-dimer interface in the aligned complexes in (**A**). Cevipabulin and compound **2** were shown in sticks. Cev: cevipabulin. cpd2: compound **2**.

